# Functional domain studies uncover novel roles for the ZTL Kelch repeat domain in clock function

**DOI:** 10.1101/2020.06.26.173096

**Authors:** Ann M. Feke, Joshua M. Gendron

## Abstract

The small LOV/F-box/Kelch family of E3 ubiquitin ligases plays an essential role in the regulation of plant circadian clocks and flowering time by sensing dusk. The family consists of three members, ZEITLUPE (ZTL), LOV KELCH PROTEIN 2 (LKP2), and FLAVIN-BINDING KELCH REPEAT F-BOX PROTEIN 1 (FKF1), which share a unique protein domain architecture allowing them to act as photoreceptors that transduce light signals via altering stability of target proteins. Despite intensive study of this protein family we still lack important knowledge about the biochemical and functional roles of the protein domains that comprise these unique photoreceptors. Here, we perform comparative analyses of transgenic lines constitutively expressing the photoreceptor LOV domain or the Kelch repeat protein-protein interaction domains of ZTL, FKF1, and LKP2. Expression of each domain alone is sufficient to disrupt circadian rhythms and flowering time, but each domain differs in the magnitude of effect. Immunoprecipitation followed by mass spectrometry with the ZTL Kelch repeat domain identified a suite of potential interacting partners. Furthermore, the ZTL Kelch repeat domain mediates interaction with the LOV domain of ZTL and the ZTL homologs LKP2 and FKF1. This suggests that the Kelch repeat domain of ZTL may mediate homo- and hetero-dimerization of the three LOV/F-box/Kelch proteins and provide added insight into the composition of the protein complexes and an additional role for the Kelch repeat domain.

## Introduction

Sensing day-night transitions is essential for proper adaptation of an organism to its environment. In plants, dusk is particularly important as it can communicate photoperiod duration and thereby the season to the plant. A multitude of biological processes depend on seasonal timing, such as reproduction, energy production, and starch biosynthesis and degradation rates that balance plant growth with night-time survival [1–4].

A small family of blue-light photoreceptors, the LOV/F-box/Kelch proteins, communicates the dusk transition to the photoperiodic flowering time pathway and the circadian clock in plants. In Arabidopsis, this family consists of three members, *ZEITLUPE* (*ZTL*), *FLAVIN-BINDING KELCH REPEAT F-BOX PROTEIN 1* (*FKF1*), and *LOV KELCH PROTEIN 2* (*LKP2*) [5–7]. These proteins act to both stabilize and de-stabilize target proteins in a light-dependent manner, thus regulating the abundance of target proteins in accordance with day/night and seasonal cycles. In order to do so, this family leverages its unique domain architecture, consisting of an N-terminal, blue-light sensing LIGHT/OXYGEN/VOLTAGE (LOV) domain, a C-terminal Kelch repeat domain, and a centrally-located F-box domain allowing it to form an E3 ubiquitin ligase that promotes ubiquitylation and subsequent proteasomal degradation of target proteins [8,9].

Despite the similarities in primary amino acid sequence between these three proteins, they display a complex pattern of genetic redundancy. For example, *ZTL* plays a major role in the regulation of the circadian clock and has only a minor role in the regulation of flowering time, while *FKF1* is an essential regulator of photoperiodic flowering that has minimal impact on the circadian clock [5,7,10–13]. *LKP2* is redundant with *ZTL* and *FKF1*, as single knockout mutations in this gene lead to minimal phenotypic consequences but exaggerate *ZTL* and *FKF1* mutant phenotypes [11,12,14]. However, when LKP2 is expressed at high levels it can cause the clock to be arrhythmic, indicating its role in clock function [6].

In order to fully understand the overlapping and distinct functions of this important gene family, intense research has begun to investigate the structures and biochemical functions of the LOV and Kelch repeat domains [9,11,22–24,14–21]. The N-terminal LOV domain is a blue light photoreceptor that is critical for the regulation of ZTL, LKP2, and FKF1 function. Regulation occurs through light-dependent interaction with the regulatory protein GIGANTEA (GI) which can promote or inhibit E3 ligase activity depending on the target protein [21,25,26]. GI binding is required for the E3 ubiquitin ligase function of FKF1, and it restricts this activity the light period of the day. GI inhibits the E3 ubiquitin ligase activity of ZTL, restricting it to the dark period [21,25,26]. Interestingly, GI is also required for the stability of the ZTL protein during the day and performs this action by recruiting the deubiquitinating enzymes UBP12 and UBP13 and acting as a co-chaperone with HSP90 [21,27,28].

In addition to its role in promoting protein interactions with regulatory proteins such as GI, the LOV domain is also involved in directly binding substrate proteins that are stabilized or destabilized. For example, the FKF1 LOV domain interacts with the floral activator CONSTANS (CO) in a GI-dependent manner, stabilizing CO in the light [19]. In contrast, the ZTL LOV domain interacts with the core clock repressors TOC1, PRR5, and CHE, promoting their degradation in the dark [13,29–31].

Adjacent to the LOV domain is a typical F-box domain and at the C-terminus of the protein is a Kelch repeat type protein-protein interaction domain. The F-box domain is a critical component of the SKP1/CULLIN/F-BOX (SCF) multi-subunit E3 ubiquitin ligase that is required for interaction with the other components of this complex [32–34]. We have previously demonstrated that the F-box domain is required for proper function of the LOV/ F-box/Kelch family of proteins, as the expression of “decoy” versions of *ZTL* and *FKF1* that lack the F-box domain mimics published loss-of-function mutant phenotypes [13]. One role of the Kelch repeat domain is to promote interactions with substrates that are ubiquitylated. This information comes from studies of FKF1, where the Kelch repeat domain binds to and promotes the degradation of the floral repressors called the CYCLING DOF FACTORs (CDFs) [24]. Interestingly, ubiquitylation of these substrates relies on the interaction between FKF1 and GI [24,26].

In contrast, the ZTL Kelch repeat domain has not been demonstrated to interact with known ZTL substrates [13,29–31]. However, the Kelch repeat domain is presumably important for ZTL function, as mutations in the Kelch repeat domain ablate ZTL function [10,20,30,35]. Additionally, expression of a truncated form of *ZTL* which contains only the LOV and F-box domains lengthens period similarly to a *ztl* loss-of-function mutant rather than shortening period as is observed in plants overexpressing full-length *ZTL [7,10,22]*. Together these data show that the Kelch repeat domain is important for ZTL’s role in clock function, but its exact biochemical role remains unknown.

In this study, we investigate the genetic and biochemical functions of the ZTL Kelch repeat domain. By performing comparative genetic analyses of plants overexpressing the *ZTL, LKP2*, and *FKF1* LOV and Kelch repeat domains, we demonstrate that both the LOV and Kelch repeat domains are independently sufficient to disrupt the circadian clock and flowering time when expressed in plants. We then focus further studies on the biochemical role of the ZTL Kelch repeat domain using immunoprecipitation followed by mass spectrometry to identify a list of putative protein interacting partners. We find that FKF1 and LKP2 as well as the native ZTL protein are part of a complex with the ZTL Kelch repeat domain. Using yeast-two-hybrid analyses, we determine that the Kelch repeat domain interacts directly with the LOV domain of ZTL suggesting two possibilities: an antiparallel conformation for intermolecular homodimers or intramolecular interaction between the two domains. Our results suggest that a biochemical role of the ZTL Kelch repeat domain may be promoting homo- or hetero-dimerization with other LOV/F-box/Kelch proteins supporting previous data providing a possible mechanism for the role of ZTL in promoting auto-ubiquitylation and also mediating stability of FKF1. Furthermore, our genetic analysis suggests that the Kelch repeat domain may modulate the formation of higher order protein complexes that are essential for the function of LOV/F-box/Kelch proteins.

## Results

### Expression of the LOV domain of LOV/F-box/Kelch proteins disrupt the circadian clock and flowering time

We have previously shown that expressing ZTL, LKP2, and FKF1 without the F-box domain results in disruption of circadian clock function and flowering time [13]. We next wanted to explore the roles of individual LOV and Kelch repeat domains separately and determine their effects on circadian clock pacing and flowering time. To do this, we overexpressed affinity-tagged LOV and Kelch repeat domains of *FKF1, LKP2*, and *ZTL* in the *CCA1p::Luciferase* (*CCA1* promoter driving expression of firefly *Luciferase*) background, and monitored circadian clock period and flowering time. We included *CCA1p::Luciferase* plants that express *FKF1, LKP2*, and *ZTL* decoys (LOV-Kelch fusion proteins which lack the F-box domain), which we have analyzed previously [13], and wild type *CCA1p::Luciferase* parental plants, as controls. In order to compare results from experiments performed separately, we use the difference between the period or flowering time of the individual T1 transgenic and the average period or flowering time of the concurrent wild type control plants for our statistical analyses [36,37]. The data generated in these experiments is displayed in Figure 1 and Tables 1 and 2. We track period and flowering time for a large number of individual T1 transgenic insertion plants allowing us to avoid potential pitfalls of following single insertion lines that may be affected by genomic insertion location.

**Table 1.** Summary of Circadian Phenotypes for *ZTL, LKP2*, and *FKF1* Decoy, LOV, and Kelch Repeat Domains.

| Domain | Subpopulation | Circadian Period Difference (Hours) |  |  |
| --- | --- | --- | --- | --- |
|  |  | ZTL (n) | LKP2 (n) | FKF1 (n) |
| Decoy | Majority | 4.5 (24)** | 1.9 (34)** | 0.6 (35)** |
|  | Minority | 0.2 (8) | -0.8 (5)* | - |
| LOV | Majority | 4.1 (25)** | 1.3 (26)** | 0.8 (36)** |
|  | Minority | 0.1 (9) | - | - |
| Kelch | Majority | 4.4 (34)** | 0.8 (19)** | 0.8 (32)** |
|  | Minority | 0.3 (3) | - | - |

The average period difference for each subpopulation of overexpression transgenics and the number of plants in each subpopulation is presented. – represents that a minority subpopulation does not exist for this domain. \* represents $p < \alpha$ Bonferroni corrected $\alpha$ of $3.8 \times 10^{-3}$ (equivalent to $p < 0.05$ ); \*\* represents $p < \alpha$ Bonferroni corrected $\alpha$ of $7.7 \times 10^{-4}$ (equivalent to $p < 0.01$ )

**Table 2.** Summary of Flowering Time Phenotypes for *ZTL, LKP2*, and *FKF1* Decoy, LOV, and Kelch Repeat Domains.

| Domain | Subpopulation | Flowering Time Difference (Days) |  |  |
| --- | --- | --- | --- | --- |
|  |  | ZTL (n) | LKP2 (n) | FKF1 (n) |
| Decoy | Majority | 2.8 (24)** | 3.8 (33)** | 6.0 (35)** |
|  | Minority | 19.9 (8)** | 29.5 (6)** | - |
| LOV | Majority | 2.6 (27)** | 4.5 (22)** | 4.1 (36)** |
|  | Minority | 16.0 (7)** | 17.0 (4)* | - |
| Kelch | Majority | 3.2 (37)** | 3.0 (19)** | 2.9 (32)** |
|  | Minority | - | - | - |

The average flowering time difference for each subpopulation of overexpression transgenics and the number of plants in each subpopulation is presented. – represents that a minority subpopulation does not exist for this domain. \* represents $p < \alpha$ Bonferroni corrected $\alpha$ of $3.8 \times 10^{-3}$ (equivalent to $p < 0.05$ ); \*\* represents $p < \alpha$ Bonferroni corrected $\alpha$ of $7.7 \times 10^{-4}$ (equivalent to $p < 0.01$ )

**Figure 1.**
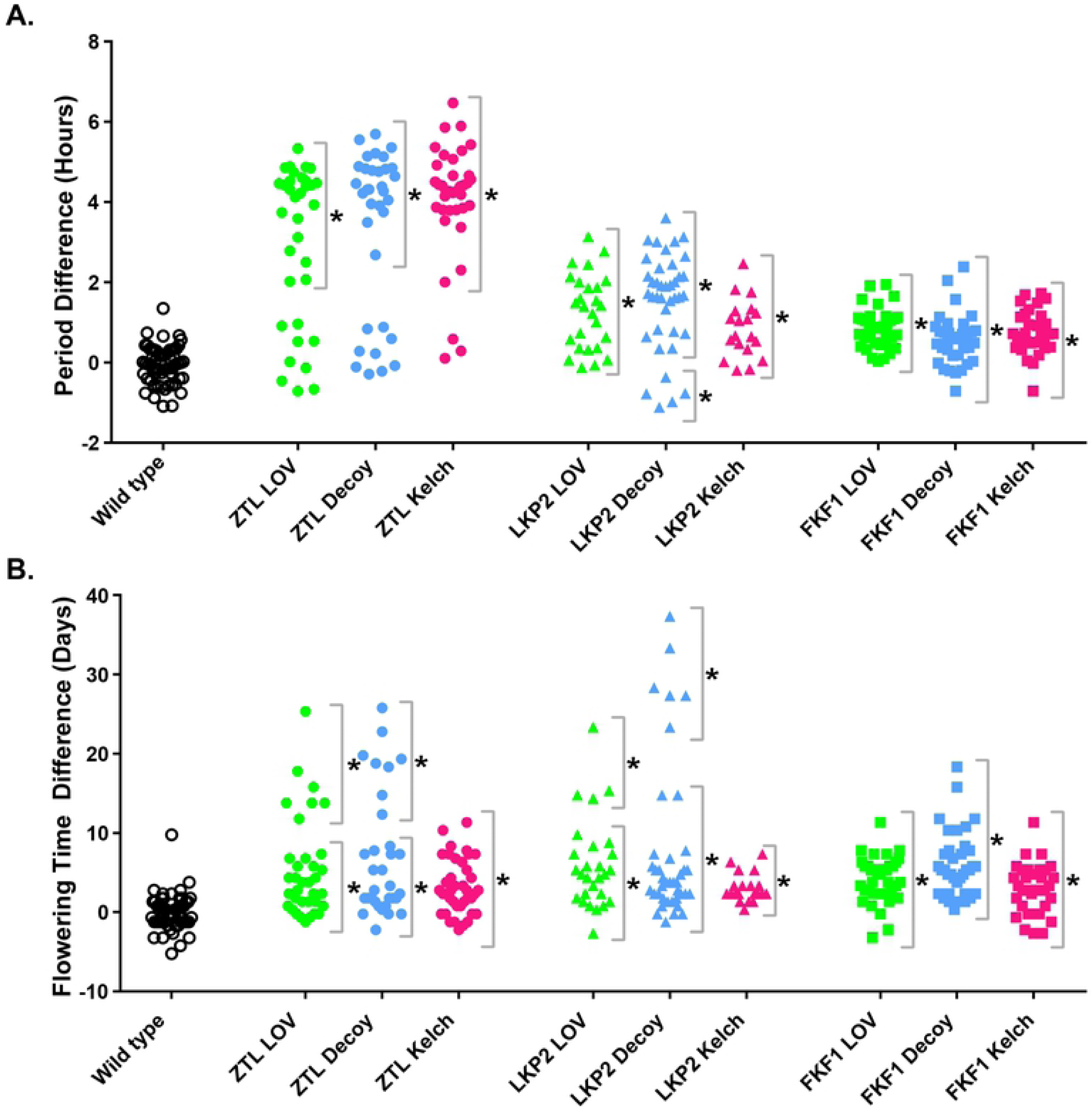
Phenotypes of Plants Expressing Domains of *ZTL, LKP2*, and *FKF1*. A) Period lengths and B) flowering time were measured for individual T1 insertion transgenics in the *CCA1p::Luciferase* background. A) Values presented are the difference between the period of the individual transgenic and the average period of the *CCA1p::Luciferase* control plants in the accompanying experiment. B) Values presented are the difference between the flowering time (as measured by the age at 1 cm inflorescence height) of the individual transgenics and the average flowering time of the *CCA1p::Luciferase* control plants in the accompanying experiment. * = significantly different from wild type with a Bonferroni-corrected α of 0.0056.

The function of the LOV domain is well defined [13,19,29–31], allowing us to predict that overexpressing the LOV domains of ZTL, FKF1, and LKP2 would be sufficient to disrupt the functions of the endogenous proteins [22]. We overexpressed affinity tagged LOV domains from *ZTL, LKP2*, and *FKF1* and monitored circadian clock period and flowering time (Fig 1A-B, green and blue points). In general, the clock and flowering phenotypes observed in plants expressing the LOV domains of ZTL, LKP2, and FKF1 are similar to the plants expressing the decoys [13]. We observed statistically significant lengthening of circadian clock period and delays in flowering time in transgenic populations expressing any of the three LOV domain and decoy constructs (Tables 1 and 2, Fig 1, green and blue points). As we have observed previously, there are phenotypic subpopulations (two separable groups of T1 transgenics from the same overexpression population) when we overexpress some decoy, LOV, or Kelch repeat domains (Fig 1A-B green and blue circles and triangles) [13]. For clarity we define the majority subpopulation as the one with the larger number of individuals, and the minority subpopulation, as the subpopulation with the smaller number of individuals.

ZTL, LKP2, and FKF1 have all been shown to regulate clock function [7,10,11,13,30]. We observed the period defect of greatest magnitude in plants expressing the *ZTL* LOV domain or *ZTL* decoy (4.1 and 4.5 hours longer, respectively), the period defect of the smallest magnitude in the plants expressing the *FKF1* LOV domain or *FKF1* decoy (0.8 and 0.6 hours longer, respectively), and a period defect of intermediate magnitude in the plants expressing the *LKP2* LOV domain or *LKP2* decoy (1.3 and 1.9 hours longer, respectively).

The relationship was inverted with regards to flowering time, with the longest delay in flowering observed in plants expressing the *FKF1* LOV domain or *FKF1* decoy (4.1 and 6.0 days, respectively), the smallest delay in flowering observed in plants expressing the *ZTL* LOV domain or *ZTL* decoy (2.6 and 2.8 days, respectively), and an intermediate delay in plants expressing the *LKP2* LOV domain or *LKP2* decoy (4.5 days and 3.8 days, respectively). Interestingly, expressing the FKF1 LOV and FKF1 decoy do not have the same effect on flowering time (4.1 days versus 6.0 days). This is consistent with the known role of the FKF1 Kelch domain in promoting the degradation of the CDF flowering time regulators [24,26].

For the minority subpopulations of plants expressing the *ZTL* LOV domain or *ZTL* decoy, we observe no statistical difference in the period from wild type, and extreme delayed flowering (16.0 days and 19.9 days late, respectively), consistent with what had previously observed in plants expressing the *ZTL* decoy (Fig 2A, blue and green circles) [13]. Interestingly, while we had not noted subpopulations in the plants expressing the *LKP2* decoy in our previous study [13], here we identify subpopulations for the plants that express the *LKP2* decoy in both period and flowering time phenotypes and subpopulations for the plants that express the *LKP2* LOV domain in the flowering time phenotypes (Figure 2B, blue and green triangles). We see a similar delay in flowering in the minority subpopulation of plants expressing the *LKP2* LOV domain (17.0 days delayed), and a more extreme delay in flowering in the minority subpopulation of plants expressing the *LKP2* decoy (29.5 days delayed). In the minority subpopulation of plants expressing the *LKP2* decoy, we also observe a small, but statistically significant, shortening of the circadian period. These results are consistent with published data that the LKP2 Kelch repeat domain may regulate CDF proteins and be important for the role of LKP2 in flowering time control [24]. In contrast, we do not observe sub-populations in the plants that express the *FKF1* LOV domain, and instead observe a positive correlation between delayed flowering and a lengthened circadian period in these plants (Figure 2C, blue and green squares), suggesting that the mechanisms through which expression of the FKF1 LOV domain lengthens period and delays flowering are linked.

**Figure 2.**
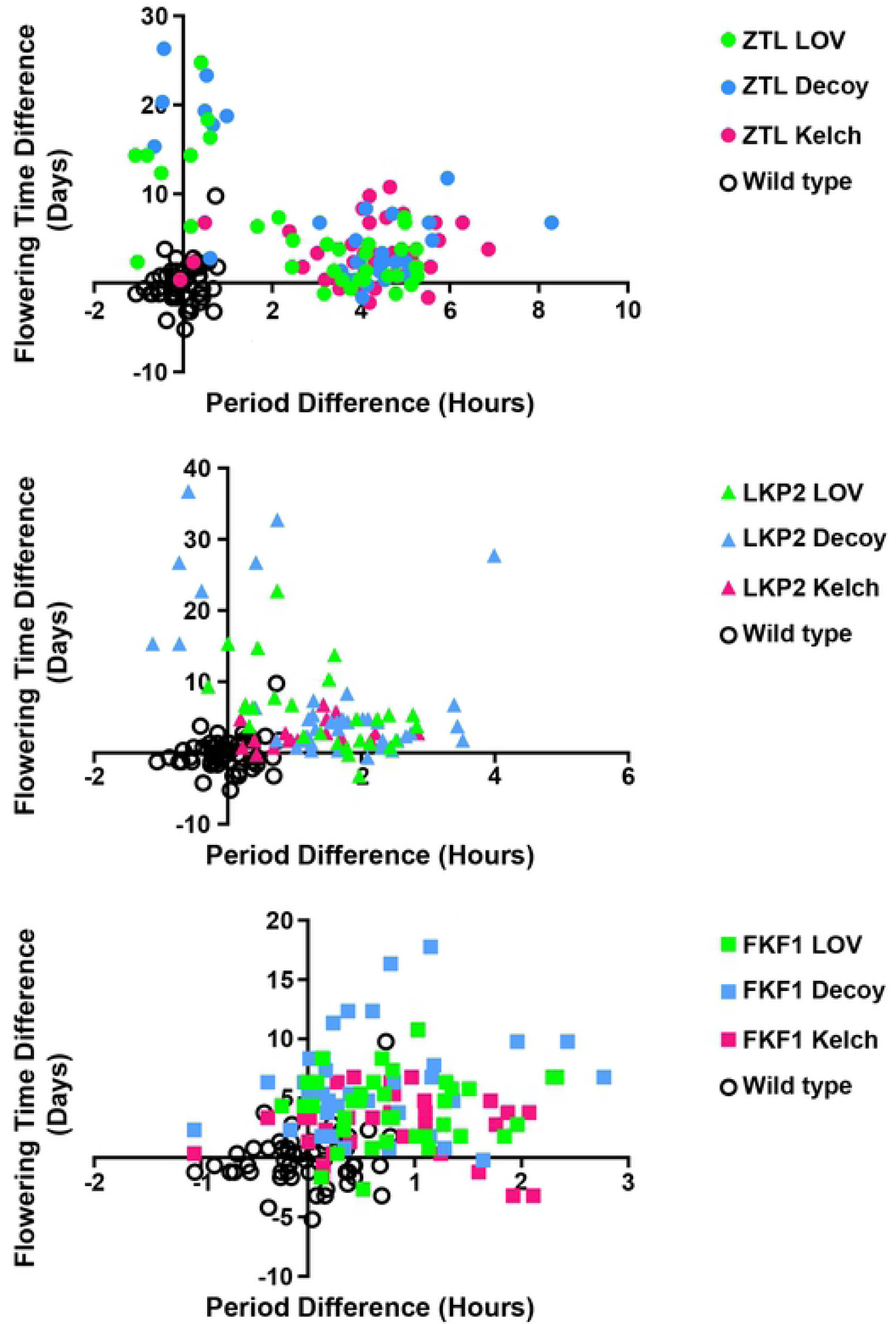
Period and Flowering Time are Anti-Correlated in the ZTL LOV, ZTL Decoy, LKP2 LOV, and LKP2 Decoy Overexpressing Plants. Data from Fig. 1 was replotted to present flowering time as a function of period for every T1 insertion plant and the *CCA1p::Luciferase* (wild type) control. A) Plants expressing domains of *ZTL* (circles); B) plants expressing domains of *LKP2* (triangles); C) plants expressing domains of *FKF1* (squares). Blue shapes: plants expressing the decoy (LOV-Kelch); Green shapes: plants expressing the LOV domain; Pink shapes: plants expressing the Kelch repeat domain; open circles: wild type plants. Note that the same wild type data is used in panels A-C.

### Expression of the Kelch repeat domains of LOV/F-box/Kelch proteins disrupt the circadian clock and flowering time control

Less is known about the Kelch repeat domain than the LOV domain of the LOV/F-box/Kelch proteins. We do know that mutations in the ZTL Kelch repeat domain cause defects in circadian clock function and the FKF1 Kelch repeat domain is needed for promoting the degradation of CDFs, demonstrating that the Kelch repeat domain is important [10,20,24,30,35]. To further explore the function of the Kelch repeat domain of the LOV/F-box/Kelch proteins we examined the effects of overexpressing affinity-tagged versions of the *ZTL, LKP2*, or *FKF1* Kelch repeat domain on the circadian clock and flowering time (Fig 1, pink points). Expressing the *ZTL, LKP2*, and *FKF1* Kelch repeat domains has similar effects on circadian period and flowering time as the majority populations of the respective decoy (Figure 1, blue and pink points, Tables 1 and 2). The most striking difference between the Kelch repeat and decoy experiments is that expressing the Kelch repeat of ZTL and LKP2 does not cause the dramatic late flowering phenotype that is caused by overexpressing the LOV domain (Fig 1, pink circles and triangles). Furthermore, we do not observe an anti-correlation between period and flowering time in the plants expressing the *ZTL* and *LKP2* Kelch repeat domain, and a weaker positive correlation in plants expressing the *FKF1* Kelch repeat domain (Figure 2,pink points). This is consistent with the idea that the LOV domain can sequester GI from the nucleus as was shown previously, but also that the Kelch repeat does not perform this function [22]. This information also demonstrates that expressing the Kelch repeat domain of the LOV/F-box/Kelch proteins can have dramatic effects on the circadian clock and flowering time confirming that this domain has an important function.

### Determining the protein interaction profile of the ZTL Kelch repeat domain

Our genetic results indicate that the ZTL Kelch repeat domain plays an important role in the regulation of the circadian clock and flowering. However, to our knowledge no biochemical function has been attributed to this domain, and it is not believed to interact with known ZTL ubiquitylation substrates or regulatory partners. We hypothesize that the ZTL Kelch repeat domain may interact with unknown substrates or regulatory partners that may help elucidate its biochemical function. Thus, we performed an immunoprecipitation followed by mass spectrometry (IP-MS) experiment in our transgenic plants constitutively expressing a HIS-FLAG tagged *ZTL* Kelch repeat domain. We collected samples from plants grown in 12 hours light/12 hours dark growth conditions at three hours before dusk (ZT9) and three hours after dusk (ZT15) (S1 Table). As controls, we included wild-type *Col-0* plants, which do not express the HIS-FLAG tag and thus control for any native Arabidopsis proteins which interact with the beads, and plants which express HIS-FLAG tagged *GFP* and thus control for any proteins which interact with the HIS-FLAG tag itself. Using this approach, we were able to identify 159 and 129 ZTL peptides from the Kelch repeat domain at ZT9 and ZT15, respectively suggesting that we were effectively immunoprecipitating the appropriate protein domain.

In our previous IP-MS studies using the ZTL decoy, we were able to identify peptides corresponding to the ZTL substrates TOC1, PRR5, and CHE and the regulatory proteins GI, UBP12, UBP13, and HSP90 [13]. While interaction studies in yeast have suggested that the Kelch repeat domain is not involved in these interactions [13,28–31], it remains possible that the ZTL Kelch repeat domain could interact with known interacting partners *in planta*. Thus, we searched our IP-MS results for peptides corresponding to the known ZTL interactors. We were unable to identify peptides corresponding to the majority of the known ZTL substrates and interacting partners (S2 Table). The only characterized interacting partners for which we were able to identify peptides were HSP90.1, HSP90.2, HSP90.4, and HSP90.5. However, we also identified peptides corresponding with these proteins in the controls. In order to determine whether the interactions between the ZTL Kelch repeat domain and HSP90 proteins was statistically significant, we performed SAINTexpress analysis [38,39] on our IP/MS results (S3 Table). We found that the interactions with HSP90.1 and 90.2 were statistically significant (SAINT score > 0.5 and Log Odds Score > 3), while the interactions with HSP90.4 and 90.5 were not statistically significant. The identification of a statistically significant interaction between the ZTL Kelch domain and HSP90 suggests that ZTL may be able to interact directly with HSP90 in the absence of GI, in addition to the ZTL-GI-HSP90 tri-partite complex that had been previously suggested [27,40]. However, lack of any peptides from other published ZTL interactors in our IP/MS results suggests that, consistent with previously published results [13,29–31], the ZTL Kelch domain does not promote interactions with the remaining known substrates and interacting partners of ZTL. It is possible that our assay was not sensitive enough to detect these interactions, but it is also possible that the ZTL Kelch domain plays an unknown role in the function of the protein through interaction with a unique group of protein partners.

We were unable to identify peptides corresponding to known ZTL targets and regulatory partners in our IP/MS results performed with the ZTL Kelch repeat domain. We next wanted to determine if other known clock or flowering time regulators interact with the ZTL Kelch repeat domain. We identified 640 statistically significant interacting proteins at ZT9, and 405 statically significant interacting proteins at ZT15. Of those proteins, 152 were identified at both ZT9 and ZT15 (Fig 3A). We had previously performed IP-MS analysis at these same time points using plants expressing the ZTL decoy. As the ZTL decoy contains the Kelch repeat domain, we would expect that high-confidence Kelch repeat interactors would immunoprecipitate with the ZTL decoy and Kelch repeat [13]. For this reason, we compared the statistically significant interactors of the ZTL decoy with the interactors we identified in this study (Fig 3B, S4 Table). We identified 50 proteins that interacted with both the ZTL Kelch repeat domain and ZTL decoy at ZT9, and 40 proteins that interacted with both ZTL isoforms at ZT15. Of those proteins, 15 were identified as statistically significant interactors of both isoforms at both time points (Table 3). Six of those 15 proteins were subunits of the T-complex, molecular chaperones that assist in protein folding [41]. We also identify metabolic enzymes, a component of the 26S proteasome, AUXIN RESPONSE FACTOR 8 (ARF8), and the ZTL homolog LKP2. These 15 proteins represent a small group of high-confidence interactors of the ZTL Kelch repeat domain.

**Table 3.**
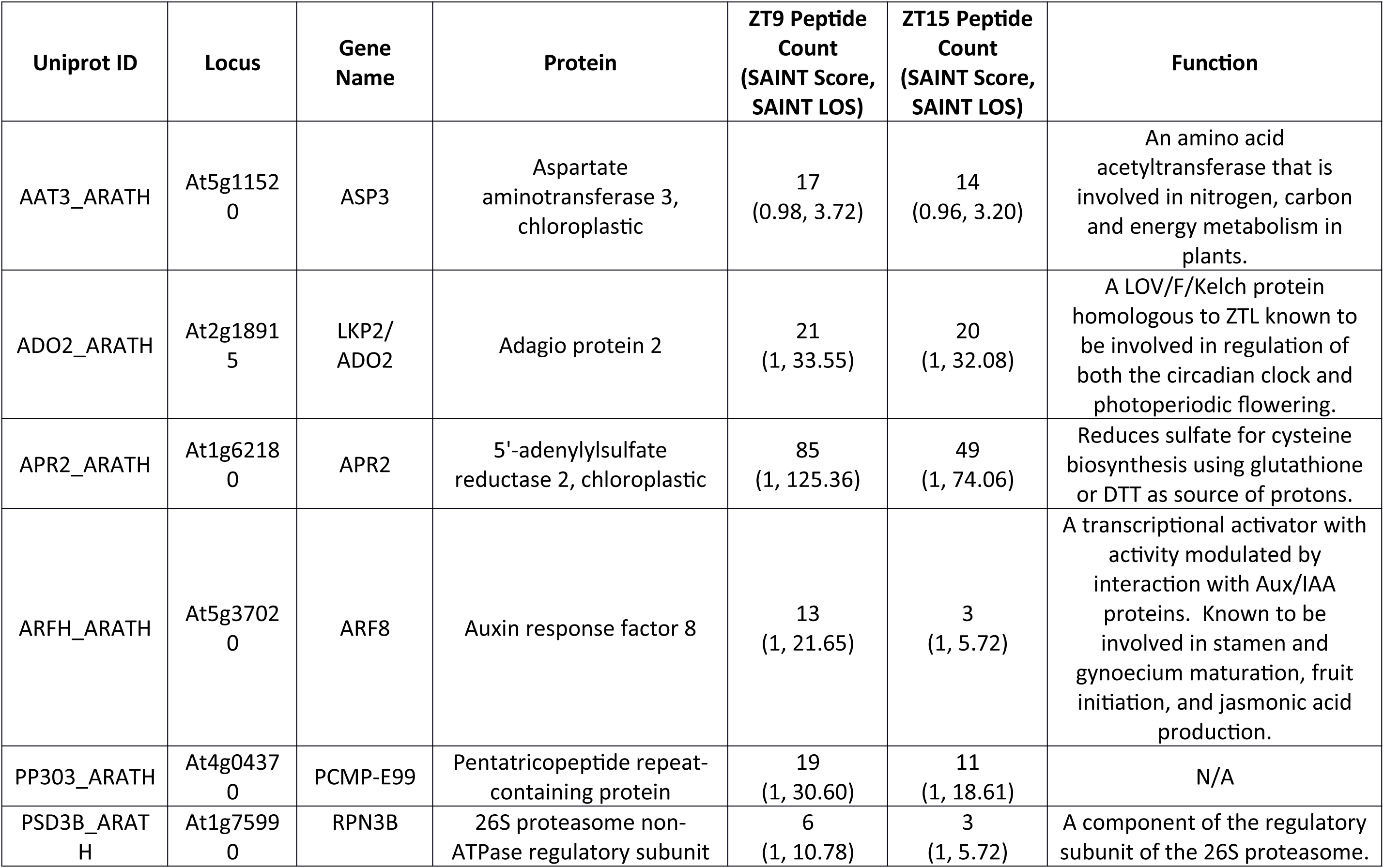
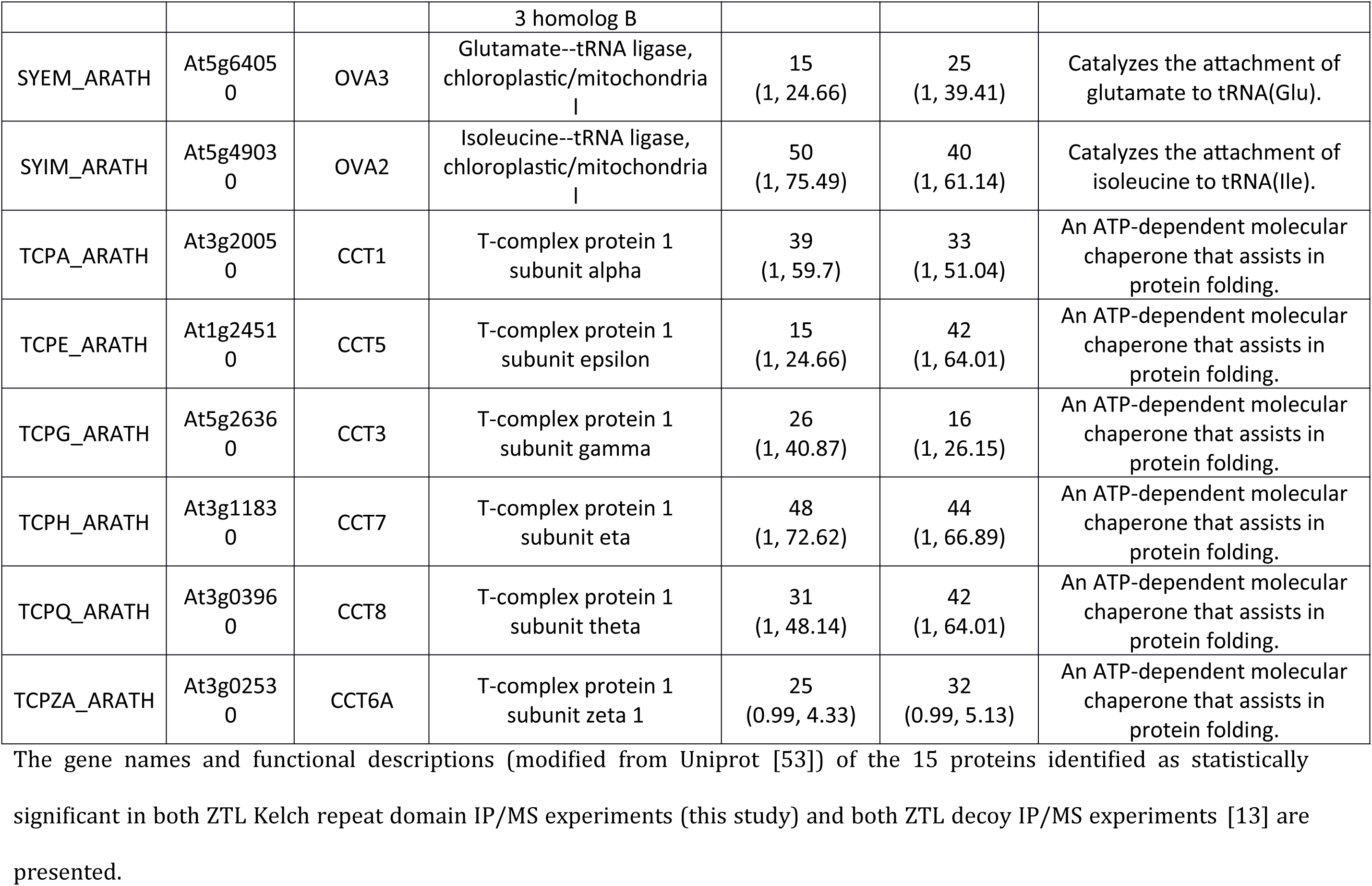
Highly Confident ZTL Kelch Repeat Domain Interactors.

**Figure 3.**
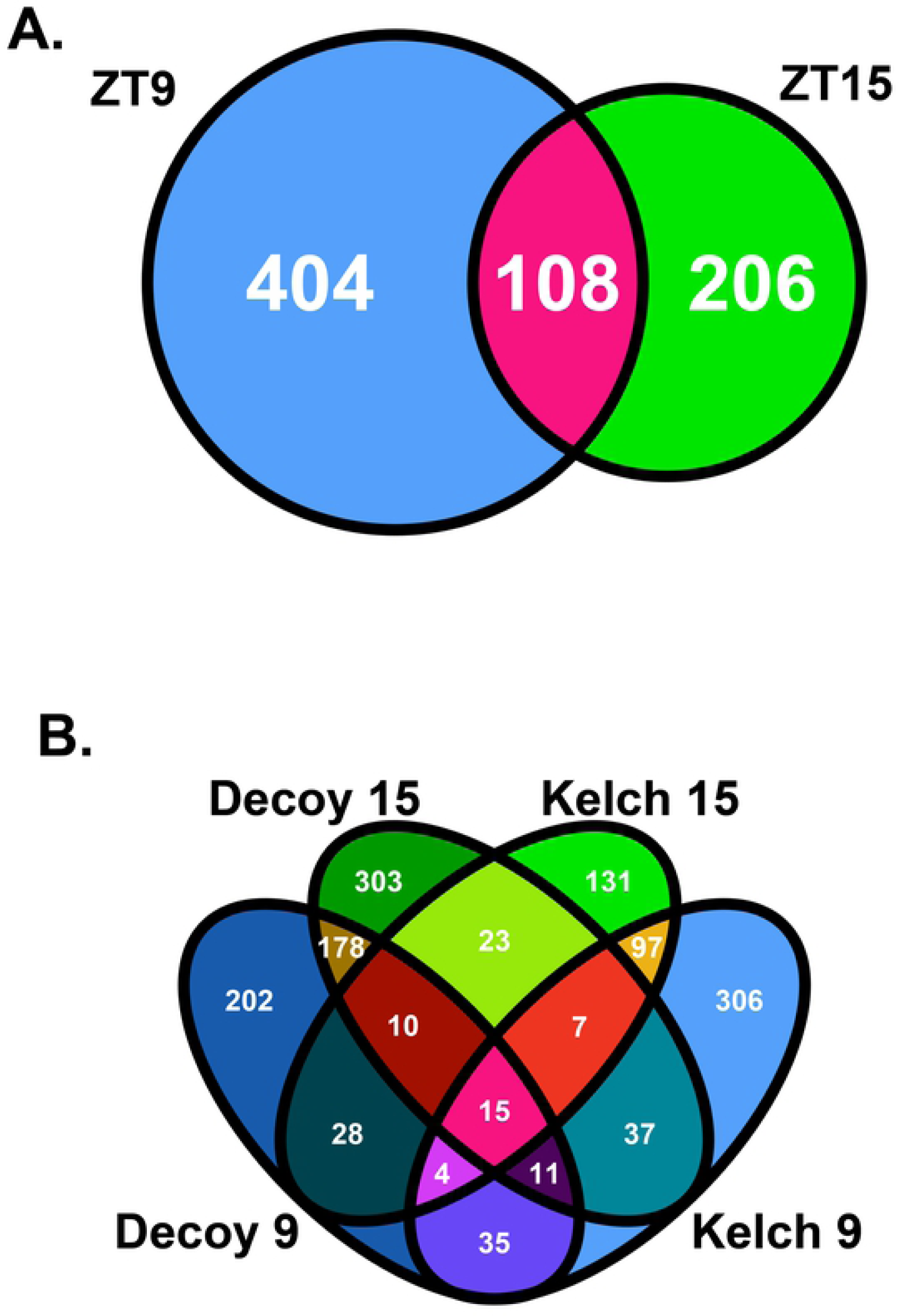
Comparison of ZTL Kelch Repeat Domain Interaction Profiles. A) A Venn diagram of the interaction profiles of the ZTL Kelch repeat domain at ZT9 and ZT15. B) A Venn diagram of the interaction profiles of the ZTL Kelch repeat domain at ZT9 and ZT15 with the ZTL decoy from [13] at ZT9 and ZT15.

We have identified a group of time and light independent ZTL Kelch repeat interacting proteins, but we also wondered if there are time dependent interactors [13,14,31]. There were 50 high-confidence interactors of both the ZTL Kelch repeat domain and the ZTL decoy at ZT9 and 40 at ZT15 (Figure 3B, S4 Table). These proteins include numerous biosynthetic enzymes, additional components of the T-complex, and, at ZT9, HSP90.1 (S4 Table). FKF1 was also identified as a statistically significant interactor of the ZTL Kelch repeat domain at ZT9 (29 peptides), but not at ZT15 (0 peptides; Fig 4A-B). This aligns well with previous reports that show the ZTL Kelch repeat domain promotes interaction with FKF1 in heterologous systems [13,14,31]. We did not observe light-dependency for the interaction between the ZTL Kelch repeat and LKP2, as equal numbers of peptides at ZT9 and ZTL15 (20 and 21 peptides, respectively) were observed (Fig 4C-D). These results confirm previous studies that suggest the ZTL Kelch repeat domain can promote heterodimerization *in planta*, but also expand on this idea and suggest that the interaction with FKF1 may be light dependent.

**Figure 4.**
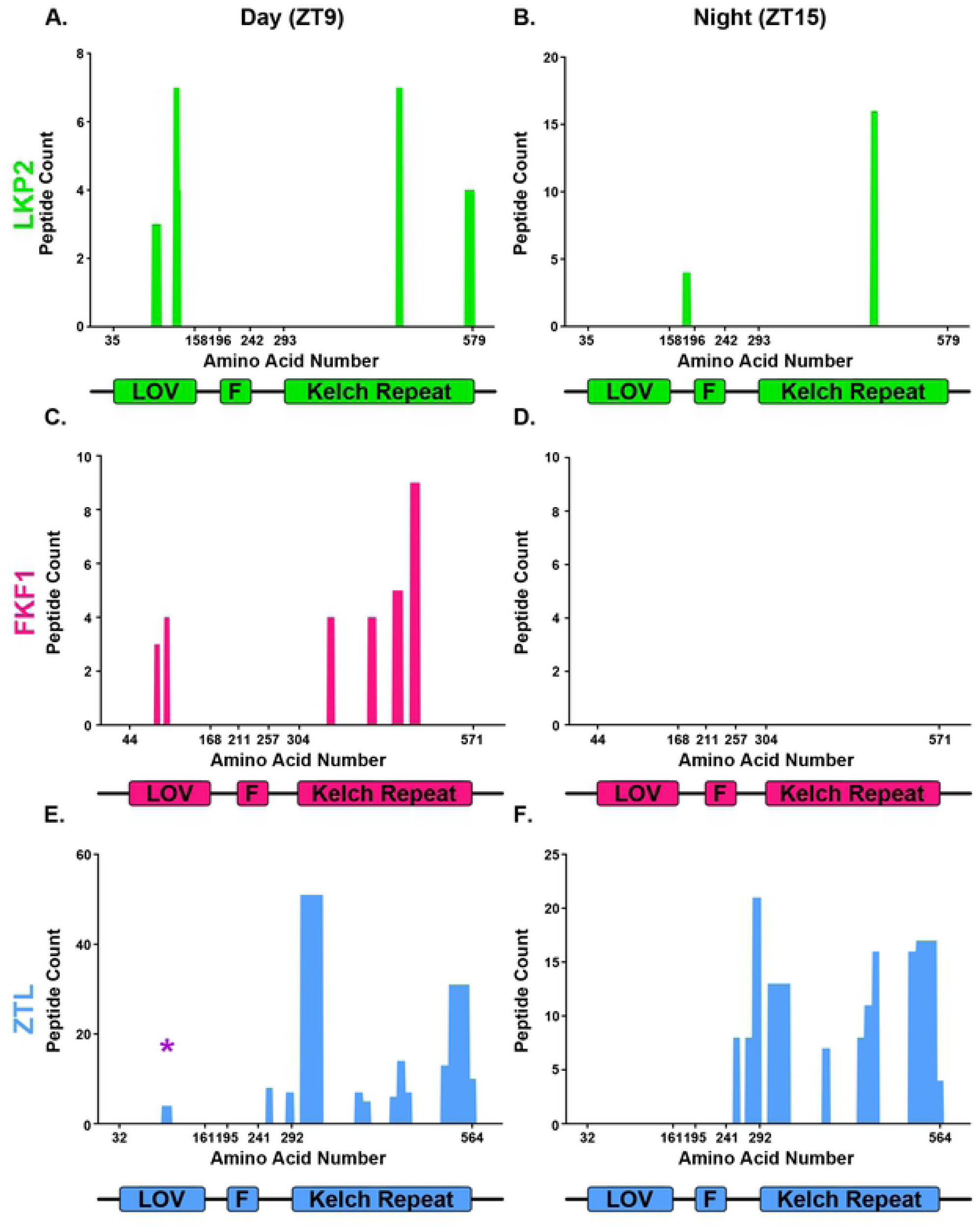
LOV-F-Kelch Family Peptides Identified in the ZTL Kelch Repeat Domain IP-MS. Peptide counts are plotted with respect to the location of the peptide within the protein sequence. Domain schematics of ZTL, LKP2 and FKF1 are included below the plots for reference, and the numbers on the x-axis represent domain boundaries. A-B) LKP2, C-D) FKF1, E-F) ZTL, A, C, E) ZTL Kelch repeat IP-MS experiment performed at ZT9; B, D, F) ZTL Kelch repeat IP-MS experiment performed at ZT15.

The ZTL decoy is capable of interacting with the native ZTL protein, suggesting that ZTL is also capable of homodimerization [13]. However, it is unclear whether the ZTL Kelch repeat domain is sufficient to drive homodimerization. In order to determine whether we identify any peptides belonging to the native ZTL protein, we aligned each ZTL peptide identified by IP-MS to the ZTL protein sequence, and mapped it to the corresponding domain (Fig 4 E-F). As expected, the majority of peptides belong to the region after the F-box domain that contains the Kelch repeat domain. However, 4 peptides (of 163 total) localized to the LOV domain (marked with a purple star). These peptides were specific for the ZTL LOV domain, and do not match any sequences in FKF1, LKP2, or the ZTL Kelch repeat domain. As with FKF1, we only observe interaction with the native ZTL protein at ZT9, however the low numbers of peptides does not exclude the possibility of interaction at ZT15 as well. This suggests that the ZTL Kelch repeat domain promotes homodimerization *in planta*.

### The ZTL Kelch repeat domain interacts with the ZTL LOV domain

Our results suggest that the ZTL Kelch repeat domain is capable of interacting with native ZTL protein, and its identification as possessing the strongest circadian effect in our phenotypic assays suggests that its expression may disrupt higher order ZTL complexes. However, it is unclear whether the interaction between the ZTL Kelch repeat and the native ZTL protein is direct, and if so, which domain of the native ZTL protein interacts with the ZTL Kelch repeat domain. Thus, we queried which domains of the ZTL protein the ZTL Kelch repeat domain is capable of interacting with by yeast-two-hybrid (Figure 5). We tested whether the ZTL Kelch repeat domain can interact with the ZTL LOV domain, with itself, with the ZTL decoy, or with the full-length ZTL protein. No interaction between the ZTL Kelch repeat domain and itself was observed. However, the ZTL LOV domain and the ZTL Kelch repeat domain did exhibit an interaction. This suggests that the interaction observed in our IP/MS results between the ZTL Kelch repeat domain and the native ZTL protein likely took place between the native ZTL LOV domain and the HIS-FLAG-tagged Kelch repeat domain. Interestingly, we did not observe interaction between the ZTL Kelch repeat domain and either the ZTL decoy or the full length ZTL protein in our yeast two hybrid experiments, which both contain the LOV domain. While the reason for this observation is currently unclear, we can posit that the presence of the Kelch in both the full length and decoy ZTL may decrease the affinity of their LOV domains for an extra-molecular Kelch repeat domain, suggesting that the LOV-Kelch interaction may typically take place intra-molecularly under natural conditions rather than inter-molecularly.

**Figure 5.**
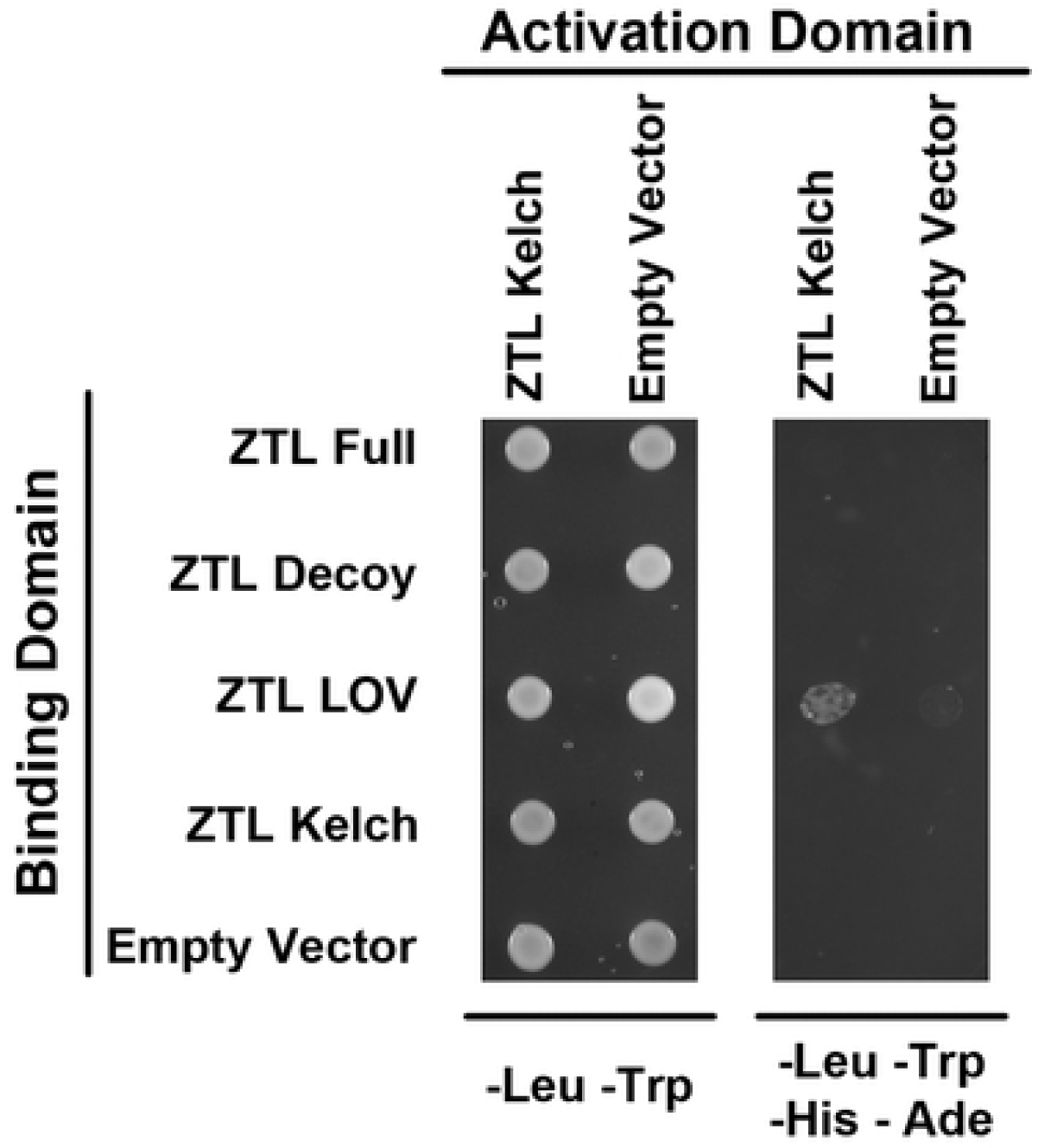
The ZTL Kelch Repeat Domain and ZTL LOV Domains Interact. Yeast-two-hybrid testing whether the activation-domain-tagged ZTL Kelch repeat domain interacts with the full ZTL protein, the ZTL decoy, the ZTL LOV domain, or the Kelch repeat domain tagged with the binding-domain.

## Discussion

### Summary

It has long been known that ZTL is an essential E3 ligase for controlling proper periodicity in the circadian clock of *Arabidopsis thaliana*. However, the precise function of each protein domain has not been fully elucidated. We had previously investigated the role of the F-box domain in this protein by characterizing plants that express “decoy” forms of ZTL and its homologs that lack the F-box domain [13]. Here, we continued this process by investigating the phenotypes of plants that express either the LOV or Kelch repeat domain of ZTL, LKP2, and FKF1, and determine that expression of either domain alone is capable of disrupting the functions of the endogenous proteins. As the ZTL Kelch repeat domain has no known function that would lead to the striking circadian phenotype we observed, we follow up with a detailed investigation of this domain. We determine the protein-protein interaction profile of the ZTL Kelch repeat domain, and identify a small suite of high-confidence interacting proteins. We also show that the ZTL Kelch repeat domain is sufficient to promote homo- and heterodimerization with the native ZTL, LKP2, and FKF1 proteins *in planta*. We find that the formation of homodimers is driven through a LOV-Kelch repeat interaction. This data suggests a new role for the Kelch repeat domain in ZTL function by promoting complex formation between ZTL and its homologs, which likely contributes to the phenotypic consequences of overexpressing this protein domain alone.

### The LOV and Kelch repeat domains contribute to ZTL, LKP2,and FKF1 function

Expression of the *ZTL* LOV domain has effects on period and flowering time which has been investigated, as has the effect of expressing the *LKP2* Kelch repeat domain on flowering time [14,22]. This study represents the first systematic and comprehensive investigation of the period and flowering time phenotypes for the LOV and Kelch repeat domains for *ZTL, FKF1*, and *LKP2*.

As a whole, we found that expression of the Kelch repeat and LOV domains of all three proteins are sufficient to produce phenotypes akin to a dominant negative effect. The period and flowering time phenotypes we observed in plants expressing the *ZTL* LOV domain and the *LKP2* Kelch repeat domain are consistent with what was observed previously [14,22]. However, to our knowledge, this is the first description of the phenotypes of plants expressing the *LKP2* LOV domain, the *FKF1* LOV domain, the *FKF1* Kelch repeat domain, or the *ZTL* Kelch repeat domain.

The ability of the ZTL LOV domain and FKF1 Kelch repeat domain to inhibit native protein function is simplest to interpret: these characterized substrate interaction domains will preferentially interact with substrates and prevent their degradation. Similar interactions likely explain the ability of the LKP2 and FKF1 LOV domains to delay period when expressed, as both domains interact with TOC1 and PRR5 [30], although the different magnitudes of these phenotypes likely represent different affinities for these substrates. Similarly, the ability of the LKP2 Kelch repeat domain to interact with the CDF proteins may cause the late flowering phenotype [24].

Not all of the observed phenotypes are as straightforward, however. For example, the delayed flowering phenotype observed in plants expressing the *FKF1* LOV domain may seem counter-intuitive. As the LOV domain of FKF1 stabilizes CO, one might expect earlier flowering [19]. However, the ability of the native FKF1 protein to degrade the CDFs is dependent on the interaction with GI [26]. Overexpressing the FKF1 LOV domain may prevent the native FKF1 protein from interacting with GI, thus preventing degradation of CDFs and leading to delayed flowering. A similar effect may explain the extremely late flowering phenotypes of the minority population of *ZTL* LOV domain and *LKP2* LOV domain expressing plants, as increased levels of the ZTL LOV domain drives GI localization towards the cytoplasm, preventing interaction between FKF1 and GI, which only occurs in the nucleus [22,26]. The absence of plants that exhibit an extremely late flowering phenotype when the *ZTL* Kelch repeat domain is expressed supports this hypothesis, as the ZTL Kelch repeat domain cannot directly interact with GI, and thus formation of the GI-ZTL or GI-FKF1 complex should be unaffected in a *ZTL*-Kelch repeat domain overexpression line [21].

To our knowledge, this study represents the first identification of a circadian defect dependent solely on the ZTL Kelch repeat domain. The identification of a large number of mutations in the Kelch repeat domain that ablate ZTL function suggests that the Kelch repeat domain is necessary [10,20,30,35]. However, previous studies have hypothesized that these mutations may affect ZTL protein function by destabilizing the structure of the entire ZTL protein [20]. Here, we have shown that the ZTL Kelch repeat domain is directly involved in circadian regulation, even in the absence of the LOV domain.

### ZTL Kelch repeat interaction profiles

In this study, we identified a large suite of proteins which may potentially interact with the ZTL Kelch repeat domain, of which 15 were identified as statistically significant interactors in both our samples here and our previous study with on the ZTL decoy [13]. Of those 15 interactors, over a third are chaperone proteins that are likely involved in the folding of the ZTL protein. Of the remainder, only one of our high confidence interactors, the ZTL homolog LKP2, is likely to play a role in circadian function. While not in our high-confidence list due to potential light-dependence, we also identify FKF1 and the native ZTL protein as putative interactors with the ZTL Kelch repeat domain.

We have noted previously that complex formation between E3 ubiquitin ligases and their homologs may be a common feature of this class of proteins, and the ability of ZTL, FKF1, and LKP2 to heterodimerize has been previously reported [13,14,31,36,37]. We demonstrate that the homodimerization takes place between the LOV and Kelch repeat domains, suggesting that the interactions between ZTL and FKF1/LKP2 may also occur in this manner. This suggests that a function of the ZTL Kelch repeat domain is to interact with the LOV domain and promote higher-order complex formation, thus modulating LOV/ F-box/Kelch protein activity.

We hypothesize that the interaction between the LOV and Kelch repeat domains of ZTL are required for its function. In support of this hypothesis, plants overexpressing a truncated form of *ZTL* containing only the LOV and F-box (ZTL LOV-F) are phenotypically indistinguishable from plants overexpressing the *ZTL* LOV domain alone [22]. If the Kelch repeat domain was dispensable for substrate ubiquitylation, one would expect that the plants overexpressing *ZTL* LOV-F would shorten period like plants overexpressing the full-length *ZTL* protein [10,22]. However, as the expression of the ZTL LOV-F protein lengthens the period, it suggests this truncated form is non-functional, suggesting that the presence ZTL Kelch repeat domain is required for proper substrate ubiquitylation.

### The ZTL LOV-Kelch repeat interaction model

We have demonstrated that the ZTL Kelch repeat domain can promote hetero- and homo-dimerization. By incorporating these interactions into models of ZTL protein function, we may begin to explain a structural conundrum of ZTL function. As F-box proteins typically have their substrate recognition domains on the C-terminus of the protein, the LOV domain is not located in a typical location for substrate ubiquitylation [32,42] and thus may be too spatially distant from the E2 conjugating enzyme to ubiquitylate LOV substrates (Fig 6A). An interaction between the LOV and Kelch repeat domains would bring the LOV domain into proximity with the E2 conjugating enzyme, and thus substrate ubiquitylation would occur (Figure 6B-C). Under this model, introduction of a truncated ZTL Kelch repeat domain would lead to the production of non-functional complexes by blocking the conformational change that brings the substrate-bound LOV domain into proximity of the E2 conjugating enzyme, potentially leading to the dominant negative phenotypes that we have observed here.

**Figure 6.**
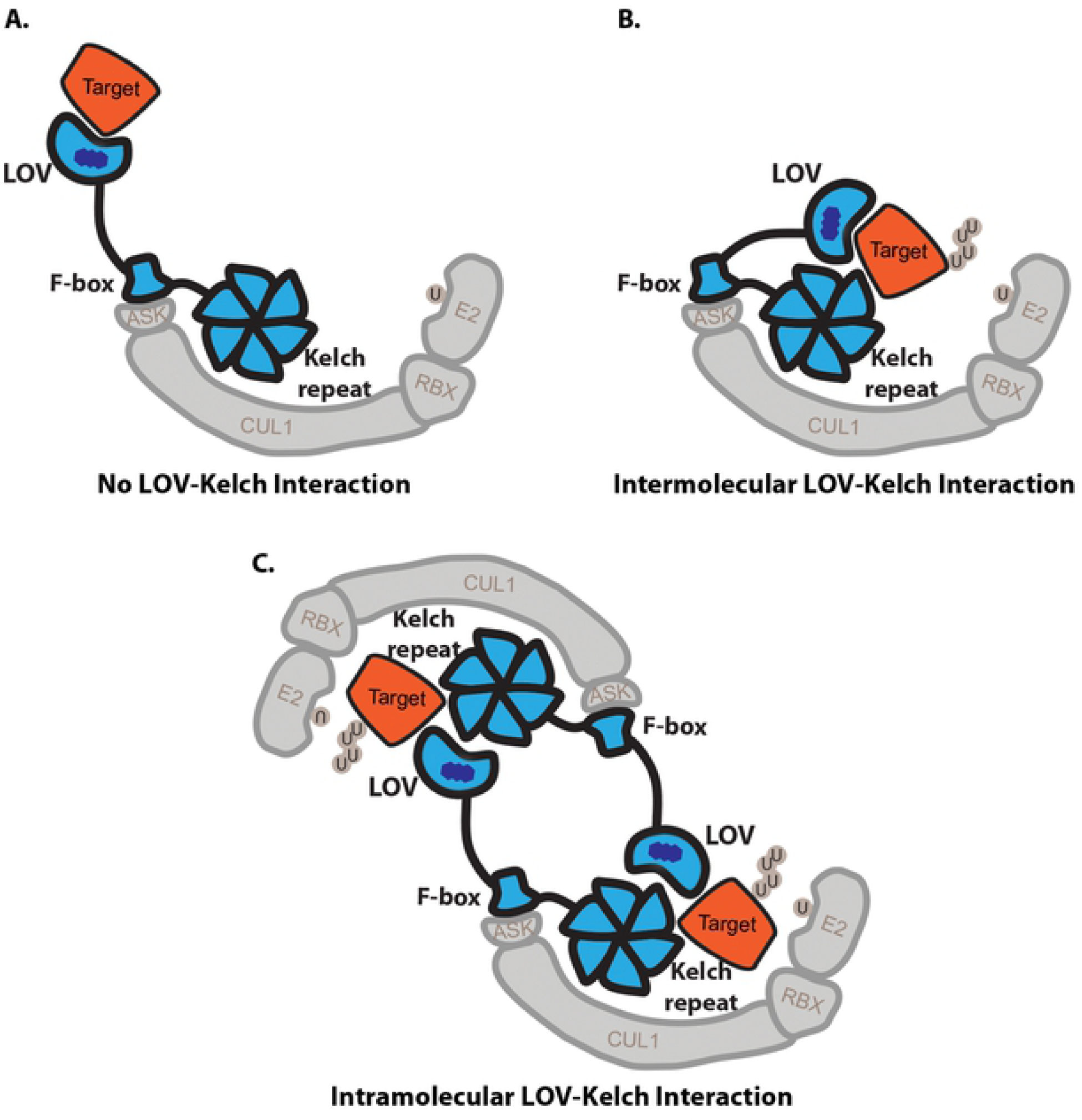
ZTL Models Demonstrate the Importance of the Kelch Repeat-LOV and Kelch Repeat-Substrate Interaction. A) Under the traditional LOV-substrate interaction without a LOV-Kelch interaction, the substrate may be too distant for ubiquitylation by a bound E2 conjugating enzyme. B-C) When these interactions are present, the substrate is brought into proximity with the E2, allowing its ubiquitylation. This interaction may either occur B) inter-molecularly, by folding the LOV domain towards the Kelch repeat domain of the same ZTL protein, or C) intra-molecularly. In the case of the intra-molecular model, two ZTL proteins share two ZTL substrates, with the LOV of one ZTL protein and the Kelch repeat of another interacting with the same substrate and with one another.

We cannot currently distinguish whether the LOV-Kelch repeat interaction occurs inter- or intra-molecularly. In the intermolecular model, the LOV domain and Kelch repeat domain of the same ZTL molecule interact with one another, folding the protein into a “closed” conformation to bring the substrates into proximity with the E2 conjugating enzyme (Figure 6B). This model is consistent with published data stating that ZTL occurs as a monomer *in planta [43]*. In the intramolecular model, two ZTL proteins align with one another in an anti-parallel fashion, and two substrate molecules are shared between the two ZTL proteins (Figure 6C). This model is more consistent with recent IP-MS data which suggests that ZTL interacts with itself to form higher-order complexes [13]. Furthermore, heterodimers of ZTL and LKP2 or FKF1 could form in the same manner as the anti-parallel ZTL homodimers. Future work will be required to distinguish between the inter- and intramolecular models of the LOV-Kelch repeat interaction.

It is interesting to note that we do not identify any peptides that correspond to GI in our ZTL Kelch repeat domain IP-MS experiments despite the observed interactions with the native FKF1, LKP2, and ZTL proteins. While this may be due to technical limitations, it may also be that the ZTL LOV-Kelch repeat interaction disrupts the LOV-GI interaction. Sequential co-immunoprecipitation experiments may prove whether the LOV-GI and LOV-Kelch repeat interactions are mutually exclusive. However, as interaction with GI inhibits ZTL E3 ligase activity [21], this suggests that the LOV-Kelch repeat conformation is the active ZTL conformation.

## Conclusions

ZTL is one of the most important E3 ligases involved in regulating the plant circadian clock, however, much is yet unknown regarding its *in vivo* structure and biochemistry. Most of the work surrounding this protein has involved the structure and function of the LOV domain, while little has covered the role of the Kelch repeat domain. Here, we have begun the process of assigning function to the Kelch repeat domain, demonstrating its importance in interactions within ZTL and its homologs. Our results illustrate the intricate interdependence of the domains of ZTL, and establish that all domains of ZTL are involved in higher order complex formation.

## Materials and methods

### Plant materials

The creation of the ZTL, LKP2, and FKF1 decoy was described previously [13]. PCR was used to amplify the LOV and Kelch repeat domains of ZTL, LKP2, and FKF1, including everything N-terminal of the F-box domain within the LOV constructs and everything C-terminal of the F-box domain within the Kelch repeat constructs, using the primers in S5 table. The amino acid numbers of the F-box domain can be found in Figure 4. PCR products were cloned into pENTR/D-TOPO vectors (Invitrogen, catalog no. K240020). The domains were then fused to FLAG and His tags at the N terminus and under the control of a cauliflower mosaic virus 35S promoter by recombination into the plant binary pDEST vector pB7-HFN [44,45] using LR recombination. The decoy constructs were transformed into Arabidopsis (*Arabidopsis thaliana*) Col-0 expressing the circadian reporter *CCA1p::Luciferase* [46] by the floral dip method [47] using *Agrobacterium tumefaciens* GV3101.

### Phenotypic analysis

Control *pCCA1∷Luciferase* and transgenic seeds were surface sterilized in 70% ethanol and 0.01% Triton X-100 for 20 minutes prior to being sown on ½ MS plates (2.15 g/L Murashige and Skoog medium, pH 5.7, Cassion Laboratories, cat#MSP01 and 0.8% bacteriological agar, AmericanBio cat# AB01185) with or without appropriate antibiotics (15 μg/mL ammonium glufosinate (Santa Cruz Biotechnology, cat# 77182-82-2)). Seeds were stratified for two days at 4 °C, then transferred to 12 hours light/12 hours dark conditions for seven days. Twenty seven-day old seedlings from each genotype were arrayed on 100 mm square ½ MS plates in a 10×10 grid, then treated with 5 mM D-luciferin (Cayman Chemical Company, cat# 115144-35-9) dissolved in 0.01% TritonX-100. Imaging was performed at 22 °C under constant 100 μmol m^-2^ s^-1^ white light provided by two LED light panels (Heliospectra L1). Hourly images were acquired for approximately six and a half days. Each hour, lights are turned off for a total of eight minutes in order to capture a 5 minute exposure on an Andor iKon-M CCD camera; lights are off two minutes prior to the exposure and remain off for one minute after the exposure is completed. After imaging is complete, the lights return to the normal lighting regime. The CCD camera was controlled using Micromanager, using the following settings: binning of 2, pre-amp gain of 2, and a 0.05 MHz readout mode [48]. Data collected between the first dawn of constant light and the dawn of the sixth day are used for analyses.

The mean intensity of each seedling at each time point was calculated using ImageJ [49]. The calculated values were imported into the Biological Rhythms Analysis Software System (BRASS) for analysis. The Fast Fourier Transform Non-linear Least Squares (FFT-NLLS) algorithm was used to calculate the period, phase, and relative amplitude from each individual seedling [50].

Following luciferase imaging, seedlings were transferred to soil (Fafard II) and grown at 22 °C in inductive 16 hours light/8 hours dark conditions with a light fluence rate of 135 μmol m^-2^ s^-1^. Plants were monitored daily for flowering status, recording the dates upon which each individual reached 1 cm inflorescence height. Each experiment was repeated twice with new independent T1 insertion transgenics in order to demonstrate repeatability. Data presented in figures and tables represents all experimental repeats, and raw values can be found in S6 table.

### Data normalization and statistical analysis

To allow for comparison across independent imaging experiments, data was normalized to the individual wild type control performed concurrently. The average value of the wild type control was calculated for every experiment, then this average was subtracted from the value of each individual T1 insertion or control wild type plant done concurrently. This normalized value was used for statistical analyses.

Welch’s t-test was used to compare each normalized T1 insertion population or subpopulation to the population of normalized control plants. In order to decrease the number of false positives caused by multiple testing, we utilized a Bonferroni corrected α as the p-value threshold. The α applied differs between experiments, and is noted throughout.

### Immunoprecipitation and mass spectrometry of plants expressing the ZTL Kelch repeat domain

Individual T1 *pB7-HFN-ZTL-Kelch* transgenics in a Col-0 background and control Col-0 and *pB7-HFC-GFP* were grown as described for phenotype analysis. Seven-day old seedlings were transferred to soil and grown under 16 hours light/8 hours dark at 22 °C for 2-3 weeks. Prior to harvest, plants were entrained to 12 hours light/12 hours dark at 22 °C for 1 week. Approximately 40 mature leaves from each background was collected and flash frozen in liquid nitrogen, such that each sample was a mixture of leaves from multiple individuals to reduce the effects of expression level fluctuations. Tissue samples were ground in liquid nitrogen using the Mixer Mill MM400 system (Retsch). Immunoprecipitation was performed as described previously [44,45,51]. Briefly, protein from 2 mL tissue powder was extracted in SII buffer (100 mM sodium phosphate pH 8.0, 150 mM NaCl, 5 mM EDTA, 0.1% Triton X-100) with cOmplete™ EDTA-free Protease Inhibitor Cocktail (Roche, cat# 11873580001), 1 mM phenylmethylsεlfonyl fluoride (PMSF), and PhosSTOP tablet (Roche, cat# 04906845001) by sonification. Anti-FLAG antibodies were cross-linked to Dynabeads^®^ M-270 Epoxy (Thermo Fisher Scientific, cat# 14311D) for immunoprecipitation. Immunoprecipitation was performed by incubation of protein extracts with beads for 1 hour at 4 °C on a rocker. Beads were washed with SII buffer three times, then twice in F2H buffer (100 mM sodium phosphate pH 8.0, 150 mM NaCl, 0.1% Triton X-100). Beads were eluted twice at 4 °C and twice at 30 °C in F2H buffer with 100 μg/mL FLAG peptide, then incubated with TALON magnetic beads (Clontech, cat# 35636) for 20 min at 4 °C, then washed twice in F2H buffer and three times in 25 mM Ammonium Bicarbonate. Samples were subjected to trypsin digestion (0.5 εg, Promega, cat# V5113) at 37 °C overnight, then vacuum dried using a SpeedVac before being dissolved in 5% formic acid/0.1% trifluoroacetic acid (TFA). Protein concentration was determined by nanodrop measurement (A260/A280)(Thermo Scientific Nanodrop 2000 UV-Vis Spectrophotometer). An aliquot of each sample was further diluted with 0.1% TFA to 0.1εg/εl and 0.5εg was injected for LC-MS/MS analysis at the Keck MS & Proteomics Resource Laboratory at Yale University.

LC-MS/MS analysis was performed on a Thermo Scientific Orbitrap Elite mass spectrometer equipped with a Waters nanoACQUITY UPLC system utilizing a binary solvent system (Buffer A: 0.1% formic acid; Buffer B: 0.1% formic acid in acetonitrile). Trapping was performed at 5εl/min, 97% Buffer A for 3 min using a Waters Symmetry® C18 180εm x 20mm trap column. Peptides were separated using an ACQUITY UPLC PST (BEH) C18 nanoACQUITY Column 1.7 εm, 75 εm x 250 mm (37°C) and eluted at 300 nl/min with the following gradient: 3% buffer B at initial conditions; 5% B at 3 minutes; 35% B at 140 minutes; 50% B at 155 minutes; 85% B at 160-165 min; then returned to initial conditions at 166 minutes. MS were acquired in the Orbitrap in profile mode over the 300-1,700 m/z range using 1 microscan, 30,000 resolution, AGC target of 1E6, and a full max ion time of 50 ms. Up to 15 MS/MS were collected per MS scan using collision induced dissociation (CID) on species with an intensity threshold of 5,000 and charge states 2 and above. Data dependent MS/MS were acquired in centroid mode in the ion trap using 1 microscan, AGC target of 2E4, full max IT of 100 ms, 2.0 m/z isolation window, and normalized collision energy of 35. Dynamic exclusion was enabled with a repeat count of 1, repeat duration of 30s, exclusion list size of 500, and exclusion duration of 60s.

The MS/MS spectra were searched by the Keck MS & Proteomics Resource Laboratory at Yale University using MASCOT [52]. Data was searched against the SwissProt_2015_11.fasta *Arabidopsis thaliana* database with oxidation set as a variable modification. The peptide mass tolerance was set to 10 ppm, the fragment mass tolerance to 0.5 Da, and the maximum number of allowable missed cleavages was set to 2.

To determine statistically significant interactors, we removed all proteins that only occurred in the controls, then performed SAINTexpress using interface available on the CRAPome website [38,39]. Proteins with a SAINT score of greater than 0.5 and a Log Odds Score of greater than 3 were considered statistically significant.

### Yeast two-hybrid assay

Yeast two-hybrid assays were performed according to the Yeast Protocol Handbook (Clontech, catalog no. P3024). Briefly, *ZTL* full length, decoy, LOV, and Kelch repeat coding sequences in pENTR/D-TOPO vectors were recombined into the pGBKT7-GW destination vector (Gateway-compatible pGBKT7 vector). This resulted in a translational fusion of the ZTL domains to the GAL4 DNA-binding domain [51]. These constructs were transformed into the yeast (*Saccharomyces cerevisiae*) Y187 strain. Similarly *ZTL* Kelch repeat coding sequences in pENTR/D-TOPO vectors were recombined into the pGADT7-GW vector (Gateway-compatible pGADT7 vector), resulting in a translational fusion to the GAL4 activation domain (Lu et al., 2010). These were transformed into the yeast AH109 strain. To test protein-protein interactions, diploid yeast was generated by yeast mating of Y187 and AH109 strains bearing pGBKT7 and pGADT7 vectors, respectively, and tested on synthetic dropout/-Leu-Trp and synthetic dropout/-Leu-Trp-His-Ade plates. The empty pGBKT7-GW and pGADT7-GW vectors were included as negative controls.

## Acknowledgements

We thank the Keck Proteomics Facility at Yale for materials and assistance with proteomics. We also thank Wei Liu, Chin-Mei Lee, Christopher Adamchek, Suyuna Eng Ren, and Catherine Chamberlin, for their technical support. We would also like to thank Sandra Pariseau for administrative support. Additionally, we would like to thank Chris Bolick, Eileen Williams, and the staff at Marsh Botanical Botanical Gardens for their support in maintaining plant growth spaces.

## Supporting Information

**Table S1. Unfiltered IP/MS results.** All results from the plants expressing the HIS-FLAG-tagged ZTL Kelch repeat domain, the plants expressing the HIS-FLAG-tagged GFP, and wild type plants are presented here.

**Table S1. Unfiltered IP/MS results.** All results from the plants expressing the HIS-FLAG-tagged ZTL Kelch repeat domain, the plants expressing the HIS-FLAG-tagged GFP, and wild type plants are presented here.

**Table S2. ZTL Kelch Repeat Domain Interaction Profile with Known ZTL Interactors**. Peptide counts and SAINT scores are presented for all published ZTL interactors.

**Table S3. SAINTexpress Results.** The results of statistically significant interactors of the ZTL Kelch repeat domain as produced by SAINTexpress.

**Table S4. Comparison of Significant Interactors Between the ZTL Kelch Repeat Domain and ZTL Decoy IP/MS Experiments.** IP/MS results from this study were compared with previously generated results from the ZTL decoy [13].

**Table S5. Primers in this Study.**

**Table S6. Source data for Figures 1-2.** Raw period and flowering time data for each individual plant presented in figure 1-2 are included.

